# Microtubule-dependent cell polarity regulates skin-resident macrophage phagocytosis and directed cell migration

**DOI:** 10.1101/2025.03.13.642867

**Authors:** Eric Peterman, Andrew Murphy, Ian A. Swinburne, Sean G. Megason, Jeffrey P. Rasmussen

## Abstract

Immune cells rapidly respond to tissue damage through dynamic properties of the cytoskeleton. How microtubules control immune cell functions during injury responses remains poorly understood. Within skin, tissue-resident macrophages known as Langerhans cells use dynamic dendrites to surveil the epidermis for damage and migrate through a densely packed epithelium to wounds. Here, we use Langerhans cells within the adult zebrafish epidermis as a model to investigate roles for microtubules in immune cell tissue surveillance, phagocytosis, and directed migration. We describe microtubule organization within Langerhans cells, and show that depolymerizing the microtubule cytoskeleton alters dendrite morphology, debris engulfment, and migration efficiency. We find that the microtubule organizing center positions adjacent to engulfed debris and that its position correlates with navigational pathfinding during directed cell migration. Stabilizing microtubules prevents Langerhans cell motility during directed cell migration by impairing navigation around cellular obstacles. Collectively, our work demonstrates requirements for microtubules in the dynamic actions of tissue-resident macrophages during epithelial surveillance and wound repair.

## INTRODUCTION

Immune cells must respond to wounds quickly to prevent pathogen invasion, clear debris, and promote tissue repair. Rapid responses require that immune cells navigate complex, 3D tissue environments that can range from loosely organized interstitial spaces to densely packed epithelia (Delgado and Lennon-Duménil, 2022). While studies from several systems have begun to reveal mechanisms of interstitial migration (Barros-Becker et al., 2017; Fernandes et al., 2020; Harada et al., 2012; Lämmermann et al., 2009; Nombela-Arrieta et al., 2007; Park et al., 2010; Tabdanov et al., 2021; Weber et al., 2013; Wilson et al., 2013; Yoo et al., 2010), navigation of immune cells within epithelia remains relatively understudied (Alraies et al., 2024; Renkawitz et al., 2019). Understanding how immune cells migrate through solid tissues is relevant to studies of diverse organ and tumor microenvironments.

The superficial location of skin makes it an excellent model for studying epithelial biology. The epidermis, the outermost layer of skin, contains tightly adherent layers of keratinocytes that form a watertight and mechanical barrier. Interspersed amongst, and confined by, epidermal keratinocytes are various immune cells that must navigate the dense microenvironment (Nestle et al., 2009). Upon skin damage, resident cells quickly mobilize to restore organ function: keratinocytes rearrange to restore the physical barrier, while immune cells migrate to the wound to clear cell debris (Peña and Martin, 2024). Our lab and others recently found that tissue-resident macrophages known as Langerhans cells migrate to sites of damage and promote skin repair (Lin et al., 2019; Peterman et al., 2024; Wang et al., 2024; Wasko et al., 2022). Langerhans cells populate the epidermis and other epithelial tissues such as oral and vaginal mucosa, surveilling tissue and acting as multimodal immune cells capable of emigration to lymph nodes and responding to tissue damage (Brand et al., 2023; Larsen et al., 1990; Lin et al., 2019; Peterman et al., 2024; Vine et al., 2024; Wang et al., 1999b; a, 2024; Wasko et al., 2022). The rapid migration of Langerhans cells to sites of tissue damage makes them an attractive model to study immune cell navigation, but how they pathfind through the confined epidermal microenvironment remains unknown.

Langerhans cells adopt a ramified morphology in homeostatic skin, and their long and dynamic dendrites facilitate skin surveillance and cell spacing (Kubo et al., 2009; Nishibu et al., 2006; Park et al., 2021). Upon skin damage, Langerhans cells modulate their dendritic morphology to accommodate cell engulfment (Peterman et al., 2024). Arborized and ramified cells such as neurons and microglia, respectively, rely on the actin and microtubule cytoskeletal networks to promote cell functionality. Work by our group and others suggest that actin and actin-regulating networks control Langerhans cell spacing, dendrite dynamics and migration in response to tissue damage (Delgado et al., 2024; Park et al., 2021; Peterman et al., 2024). Microtubules and the microtubule organizing center (MTOC) function prominently during immune cell migration through interstitial microenvironments (Renkawitz et al., 2019; Yoo et al., 2012) and previous studies implicate microtubules in Langerhans cell antigen uptake and presentation (Bacci et al., 1996; Martin et al., 2014), but how microtubules function during immune cell migration through solid epithelia remains an open question.

Due to its optical clarity, zebrafish skin is an excellent model for studying immune cell motility in a native microenvironment. Along the fish trunk, the epidermis drapes across plate-like scales, a type of dermal appendage **(Figure 1A, B)** (Aman and Parichy, 2024). Similar to other types of vertebrate skin, the adult zebrafish epidermis contains stratified keratinocytes and interspersed immune cells, including T cells and Langerhans cells **(Figure 1B)** (Dee et al., 2016; He et al., 2018; Hui et al., 2017; Lin et al., 2019; Robertson et al., 2023). Previously, we developed an *ex vivo* scale explant model that allows for the visualization of Langerhans cell behaviors with high spatiotemporal resolution and acute cytoskeletal manipulation via chemical perturbations (Guerrero et al., 2024; Peterman et al., 2023, 2024). Here, we use this skin explant system in combination with new transgenes to examine the function of microtubules and the MTOC during Langerhans cell skin surveillance and responses to tissue damage.

**Figure 1.**
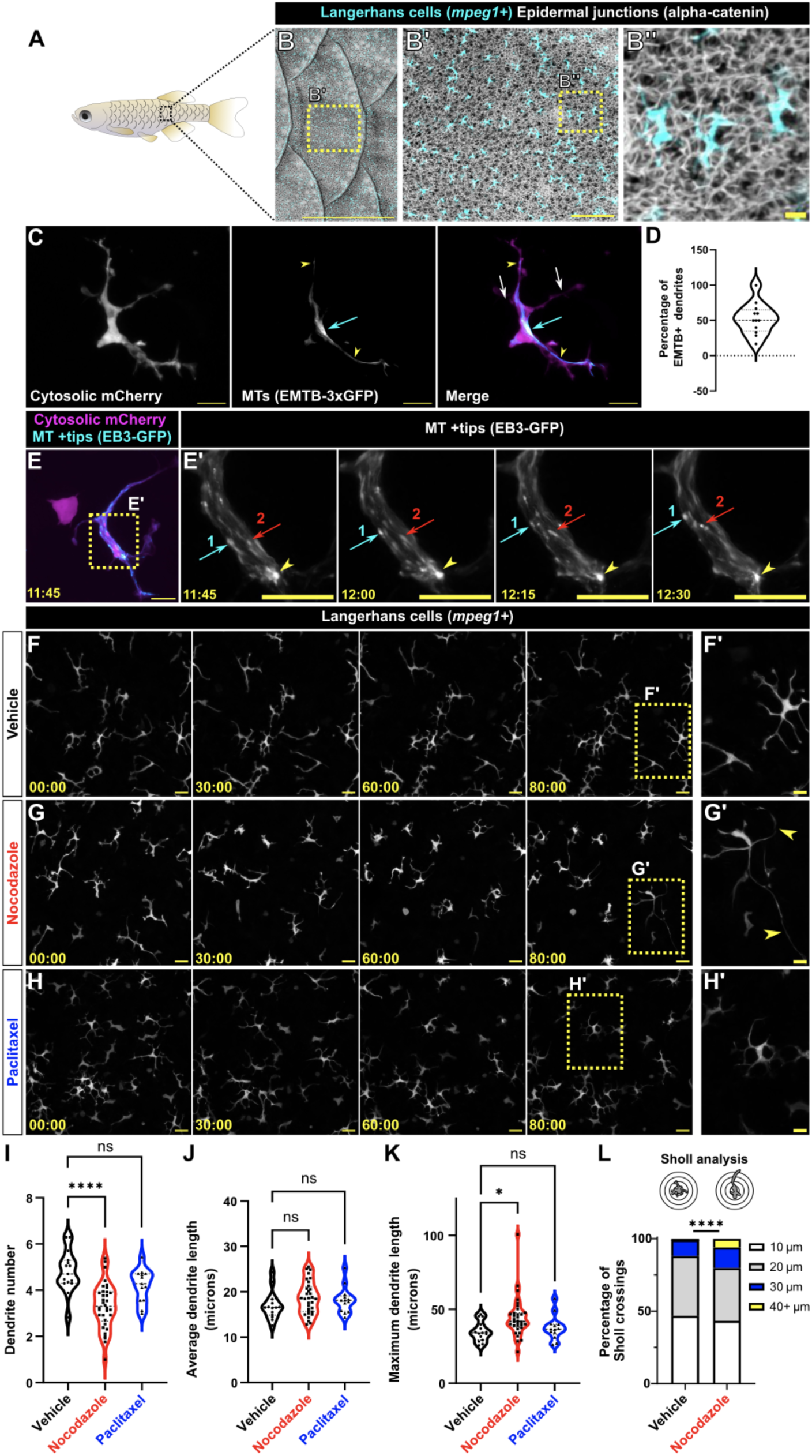
Microtubule depolymerization disrupts Langerhans cell morphology. **A.** Illustration of adult zebrafish showing overlapping scales along the trunk skin. **B.** Confocal images of adult trunk skin expressing reporters for Langerhans cells [Tg(*mpeg1:mCherry)*] and alpha-catenin [*Gt(ctnna1-Citrine)*], which labels epidermal junctions. **C.** Representative image of a Langerhans cell expressing *Tg(mpeg1.1:mCherry;mpeg1.1:EMTB-3xGFP)* showing EMTB-3xGFP+ dendrites (yellow arrowheads), EMTB-3xGFP-dendrites (white arrows), and the presumptive microtubule organizing center (cyan arrow). **D.** Violin plot showing percentage of EMTB+ dendrites *(n =* 12 cells from *N* = 5 scales). **E.** Representative still images from time-lapse confocal microscopy of a *Tg(mpeg1.1:mCherry;mpeg1.1:EB3-GFP)+* Langerhans cell showing the perinuclear focus of EB3 signal (yellow arrowhead). Yellow box in left panel denotes region magnified in the panels at right. Red and cyan arrows track individual growing microtubules. **F-H.** Representative still images from time-lapse confocal microscopy depicting effects of vehicle **(F),** nocodazole **(G)**, or paclitaxel **(H)** treatment on *Tg(mpeg1.1:YFP)*+ Langerhans cell morphology. Yellow arrowheads in **(G)** indicate elongated dendrites. **I-K.** Violin plots showing average dendrite number **(I)**, average dendrite length **(J)**, or maximum dendrite length **(K)** through 80 minutes of vehicle, nocodazole, or paclitaxel treatment. *n* = 15 cells tracked from *N* = 3 scales for vehicle control, *n =* 27 cells tracked from *N =* 5 scales for nocodazole, *n =* 13 cells tracked from *N =* 6 scales for paclitaxel. **L.** Sholl analysis of dendrite lengths in vehicle- and nocodazole-treated conditions. *n =* 698 dendrites tracked in 14 cells from *N =* 3 scales for vehicle control and n = 476 dendrites tracked in 11 cells from *N =* 3 scales for nocodazole. Statistical significance in **(I, J, K)** was determined using Mann-Whitney U test, statistical significance in **(L)** was determined using a chi-squared test with the raw number of dendrites. * = *p* < 0.05, **** = *p* < 0.0001. Timestamps denote mm:ss. Scale bars, 1 mm **(B),** 100 μm **(B’),** 10 μm **(B’’, C, E, F’-H’)**, 20 μm **(F-H).**

## RESULTS AND DISCUSSION

### Microtubule organization in Langerhans cells in homeostatic skin

The zebrafish *mpeg1.1* promoter labels diverse macrophage populations, including Langerhans cells (Ellett et al., 2011; He et al., 2018; Lin et al., 2019; Zhou et al., 2023). To visualize microtubules in Langerhans cells, we created a transgenic line in which the *mpeg1.1* promoter drives expression of the microtubule reporter EMTB-3xGFP (*Tg(mpeg1.1:EMTB-3xGFP)*) (Barros-Becker et al., 2017; Faire et al., 1999). To simultaneously visualize Langerhans cell microtubules and cytoplasm, we crossed this line to *Tg(mpeg1.1:mCherry)* (Ellett et al., 2011), which expresses cytosolic mCherry in Langerhans cells **(Figure 1C, Supplemental Video 1)**. From confocal z-stacks, we identified linear arrays of EMTB-3xGFP signal in at least one dendrite of all Langerhans cells examined (**Figure 1C, arrowheads and 1D,** range: 16.67-100% EMTB-3xGFP+ dendrites/cell; *n*=12 cells). Microtubule organization and polarity is dictated in many cell types by a single microtubule organizing center (MTOC) (Meiring et al., 2020). Previous studies have used EMTB+ foci to identify the MTOC of migrating dendritic cells (Kroll et al., 2023; Renkawitz et al., 2019). While imaging EMTB-3xGFP localization in Langerhans cells, we observed that microtubules appeared to emanate from the brightest point of signal, which localized near the nucleus **(Figure 1C, cyan arrow)**, suggesting this may represent the MTOC. As a complimentary approach to MTOC identification, we used the plus-end binding protein EB3 (Stepanova et al., 2003) to label the growing plus-ends of microtubules in Langerhans cells of *Tg(mpeg1.1:mCherry)* adult zebrafish. We found that EB3-GFP+ comets grew from a bright, perinuclear EB3-GFP+ focus **(Figure 1E; Supplemental Video 2)**. Based on these combined data, we concluded that Langerhans cells have a single, perinuclear MTOC, and that microtubules are present in Langerhans cell dendrites.

To test if microtubules are required to maintain the dendritic morphology of Langerhans cells, we explanted skin and administered nocodazole to depolymerize microtubules **(Figure 1F,G; Supplemental Video 3)**. We first confirmed that nocodazole treatment resulted in loss of EMTB-3xGFP signal **(Supplemental Figure 1)**, indicating our reporter faithfully labels microtubules and that nocodazole depolymerized microtubules in our explant system. We next analyzed Langerhans cell morphology using the cytosolic reporter *Tg(mpeg1.1:YFP)* (Roca and Ramakrishnan, 2013) and found that nocodazole treatment significantly decreased Langerhans cell dendrite number compared to vehicle-treated controls **(Figure 1I).** Although average dendrite length was only modestly increased in nocodazole-treated cells **(Figure 1J),** the maximum dendrite length increased significantly **(Figure 1K).** As an alternative measure of dendrite morphology, we used Sholl analysis to measure the number of dendrites crossing concentric circles in increasing 10 micron intervals. Our analysis revealed that microtubule depolymerization significantly increased the range of dendrite lengths **(Figure 1L).** To examine how microtubule stabilization affected Langerhans cell morphology, we also treated scale explants with paclitaxel, a microtubule-stabilizing drug (Schiff and Horwitz, 1980; Schiff et al., 1979). Treatment of scale explants with paclitaxel did not alter dendrite number or length **(Figure 1H-K).** These data indicate that acute microtubule depolymerization, but not stabilization, alters Langerhans cell dendrite number and length in steady-state conditions.

### The perinuclear microtubule organizing center rapidly localizes to sites of debris engulfment

We next questioned if the microtubule cytoskeleton was important for debris engulfment by Langerhans cells. We began by measuring MTOC motility during steady-state conditions and observed that the brightest EMTB-3xGFP focus per cell (hereafter referred to as the MTOC) moved an average of 1.39 microns/minute in the absence of stimulus **(Figure 2A,B; Supplemental Video 1).** Next, we tracked Langerhans cell engulfment of laser-damaged single keratinocytes as previously described **(Figure 2C)** (Guerrero et al., 2024; Peterman et al., 2024). Consistent with observations in microglia (Möller et al., 2022), we found that the Langerhans cell MTOC relocalized to the site of debris contact after laser ablation **(Figure 2D,E; Supplemental Video 4).** When we quantified the distance that the MTOC traveled 5 minutes prior to ablation and 5 minutes prior to engulfment, we found that the MTOC traveled further distances following Langerhans cell activation **(Figure 2F)**. Additionally, we found that MTOC speed was faster after laser ablation (average of 1.83 microns/minutes) compared to pre-ablation (average of 1.26 microns/minute) **(Figure 2G)**. Altogether, these data show that Langerhans cell MTOC motility increases after nearby cellular damage and that the MTOC relocalizes proximal to debris during engulfment.

**Figure 2.**
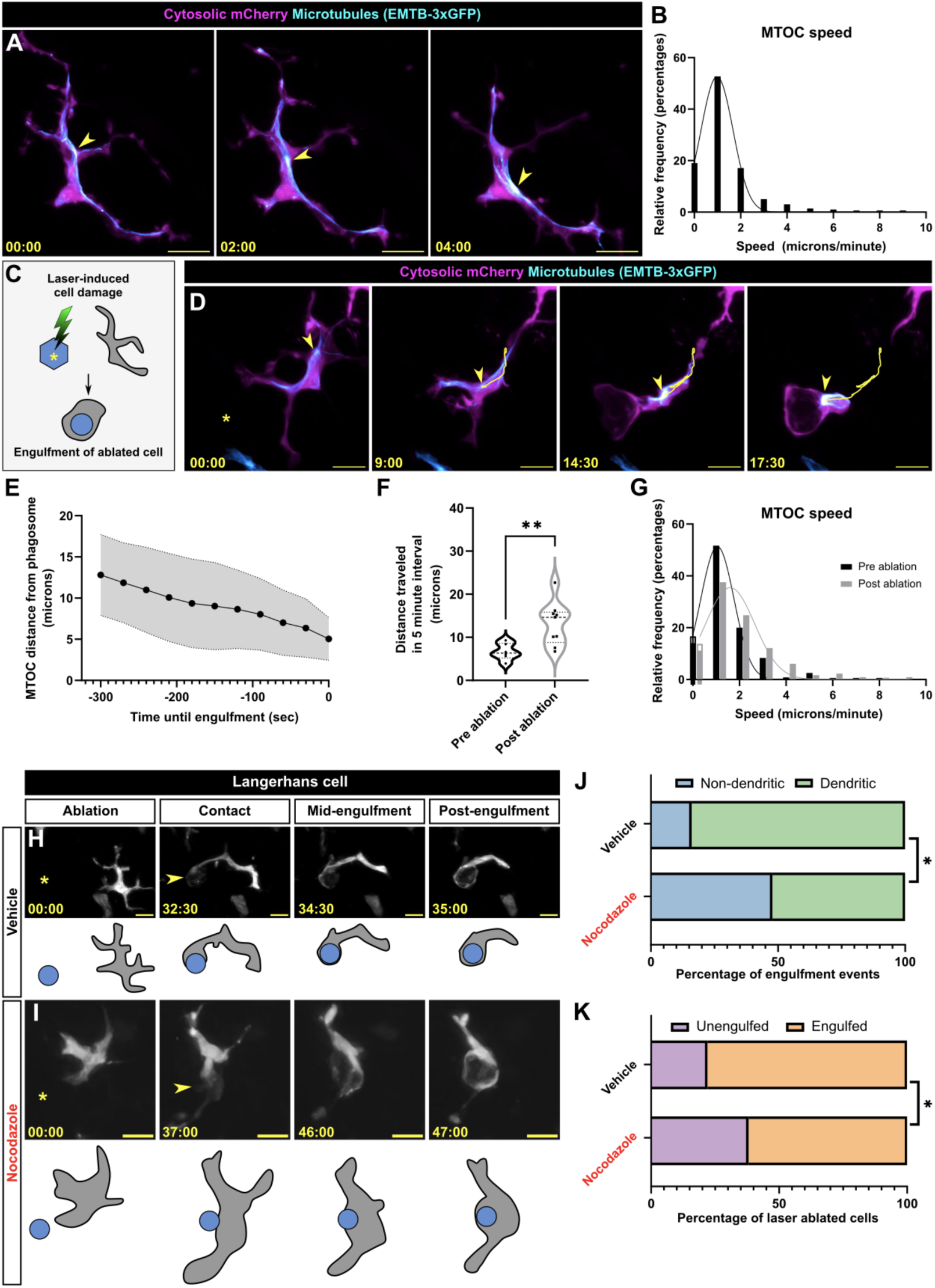
MTOC dynamics and microtubule requirements during debris engulfment. **A.** Representative still images from time-lapse confocal microscopy of a *Tg(mpeg1.1:mCherry;mpeg1.1:EMTB-3xGFP)+* Langerhans cell showing MTOC (yellow arrowhead) movement in steady-state conditions. **B.** Histogram showing the frequency distribution of MTOC speed in steady-state conditions, *n =* 10 cells tracked from *N =* 5 scales. **C.** Schematic of laser-induced keratinocyte (blue) damage and subsequent Langerhans cell engulfment. **D.** Representative still images from time-lapse confocal microscopy showing MTOC (yellow arrowhead) motility preceding engulfment of debris generated by keratinocyte laser ablation. Yellow trace indicates the MTOC track over time. Yellow asterisk indicates the site of laser ablation. **E.** Quantification of the distance between the MTOC and phagosome in the 5 minutes prior to debris engulfment, *n =* 9 cells tracked from *N =* 4 scales. Gray shading represents a 95% confidence interval. **F.** Violin plot of the total distance traveled by the MTOC 5 minutes prior to keratinocyte laser ablation and 5 minutes prior to keratinocyte engulfment, *n =* 6 cells tracked in pre ablation, *n =* 9 cells tracked in post ablation from *N =* 4 scales. **G.** Histogram of the frequency distribution of MTOC speed 5 minutes prior to keratinocyte laser ablation and 5 minutes prior to keratinocyte engulfment, *n =* 6 cells tracked in pre ablation, *n =* 9 cells tracked in post ablation from *N =* 4 scales. **H,I.** Representative still images from time-lapse confocal microscopy of vehicle-**(H)** or nocodazole-treated **(I)** *Tg(mpeg1.1:YFP)+* Langerhans cells showing engulfment of debris. Yellow asterisk indicates the site of laser ablation. Yellow arrowhead indicates site of first contact between the Langerhans cell and laser-damaged cell. **J.** Quantification of engulfment modality used to engulf debris, *n =* 25 engulfment events tracked from *N =* 3 individual experiments for vehicle control, *n =* 23 ablation events tracked from *N =* 3 individual experiments for nocodazole. **K.** Quantification of successful keratinocyte debris engulfment following addition of vehicle or nocodazole, *n =* 30 ablated cells tracked from *N =* 3 individual experiments for vehicle control, *n =* 26 ablated cells tracked from *N =* 3 individual experiments for nocodazole. Statistical significance in **(F)** was determined using a Mann-Whitney U test. A Kolmogorov-Smirnov test was used in **(G)** and revealed no significant difference. Fisher’s exact test was used to determine significance in **(J,K)** by using the raw counts of engulfment events. Traces in frequency distribution graphs **(B)** and **(G)** are Gaussian fits. * = *p* < 0.05, ** = *p* < 0.01. Timestamps denote mm:ss. Scale bars, 10 μm **(A,D,H,I)**.

### Dendrite-mediated engulfment of debris requires the microtubule cytoskeleton

Our results indicate that microtubule depolymerization alters Langerhans cell dendrite morphology and that the MTOC polarizes towards cell debris after laser ablation. This led us to hypothesize that microtubules may be required for debris engulfment. To test this, we used our laser ablation paradigm in combination with vehicle or nocodazole treatment and tracked debris engulfment. In vehicle-treated conditions, we observed two modes of debris engulfment by Langerhans cells. The predominant modality involved Langerhans cells contacting debris with a single, long dendrite, while simultaneously retracting distal dendrites prior to debris engulfment **(Figure 2H,J**; 84% of events**; Supplemental Video 5)**. Alternatively, Langerhans cells adopted a non-dendritic modality in which the entire cell migrated and repositioned itself next to the debris, which was then engulfed at the cell body **(Figure 2J**; 16% of events). By contrast, in nocodazole-treated conditions, Langerhans cells exhibited increased usage of the motility-based engulfment modality **(Figure 2I,J)** and a decrease in the ability to engulf debris **(Figure 2K)**. These results suggest that microtubules are partly responsible for dendrite-mediated engulfment and overall engulfment success of damaged keratinocytes.

### Efficient directed cell migration requires the microtubule cytoskeleton

In addition to their function in regulating phagocytosis, microtubules and the MTOC play pivotal roles in immune cell migration (Etienne-Manneville, 2013). Previous work demonstrated that murine and zebrafish Langerhans cells migrate to tissue wounds generated by punch biopsy or mechanical scratch, respectively (Wasko et al., 2022; Peterman et al., 2024; Lin et al., 2019), but the underlying mechanisms that promote migration remain unknown. To ask if Langerhans cells require microtubules for directional migration, we pretreated explanted scales with vehicle or nocodazole, introduced an epidermal scratch, and imaged cell migration over 3 hours **(Figure 3A,B; Supplemental Video 6)**. Compared to vehicle-treated controls, nocodazole-treated cells migrated with a significantly larger total distance traveled and displacement **(Figure 3C,D)**. As a measure of the efficiency of migration, we calculated the meandering index (defined as the ratio of displacement to total distance traveled) and found that Langerhans cells had a smaller meandering index, indicating the cells did not migrate as directly toward the wound **(Figure 3E)**. Confirming this, we observed significantly fewer nocodazole-treated cells in the wound margin **(Figure 3F)**. Altogether, these data indicate that microtubule depolymerization causes increased Langerhans cell motility, but with less directionality, leading to reduced cell accumulation at sites of tissue damage.

**Figure 3.**
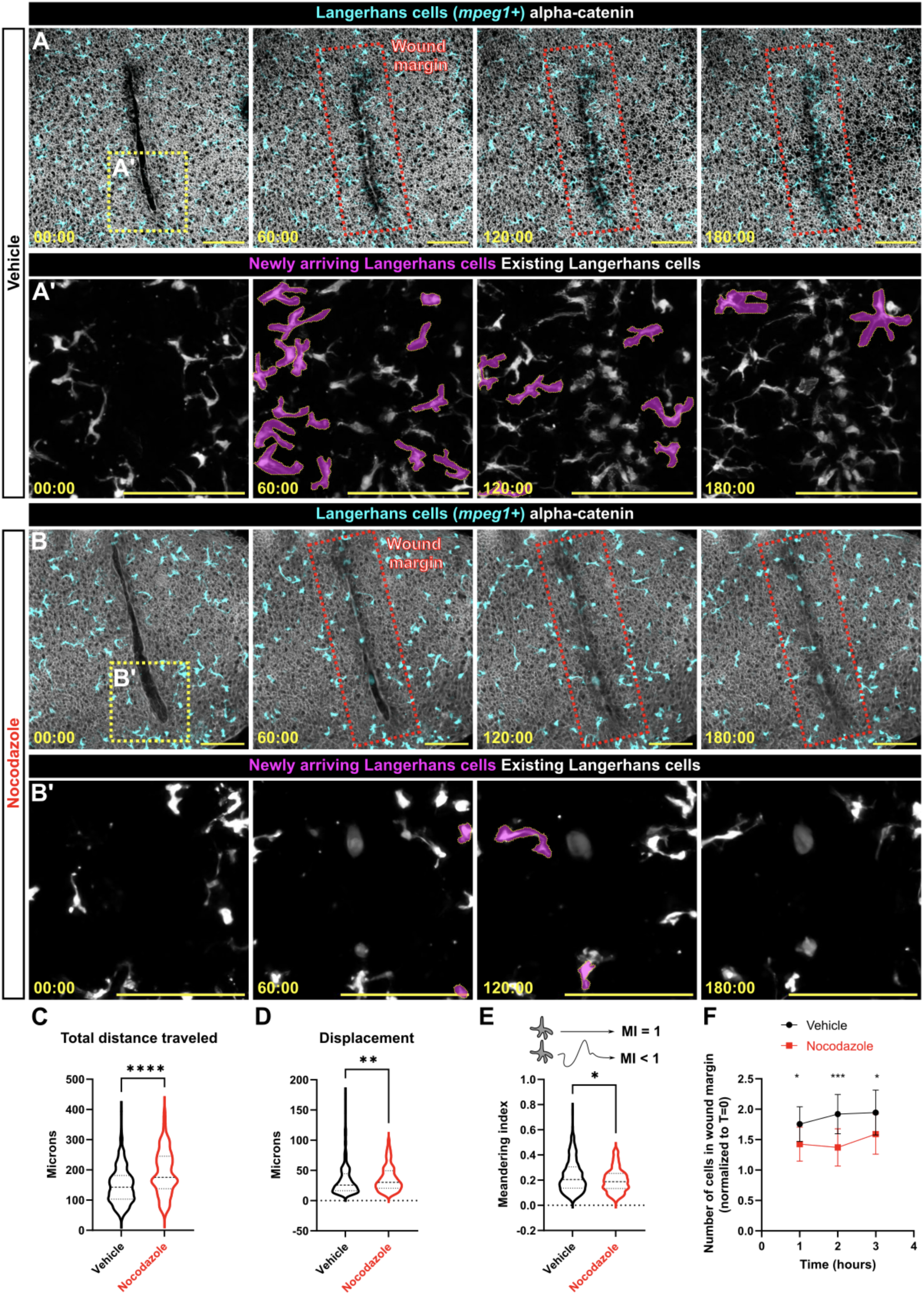
Langerhans cell migration to scratch wounds requires microtubules. **A,B.** Langerhans cell migration to epidermal scratch in vehicle-**(A)** or nocodazole-treated **(B)** conditions. Yellow boxes in first frames of **(A, B)** indicate insets in A’, and B’, red boxes in frames 2-4 indicate the wound margin ROI corresponding to cell counts in **(F).** Newly arriving Langerhans cells are pseudocolored magenta in **A’, B’. C-E.** Violin plots showing total distance traveled **(C)**, displacement **(D)**, or the meandering index **(E)** of *mpeg1+* cells over 3 hours in response to epidermal scratch in vehicle- or nocodazole-treated conditions, *n =* 325 cells from *N =* 11 skin explants tracked in vehicle conditions, *n =* 181 cells from *N =* 10 skin explants tracked in nocodazole conditions. **F.** Quantification of the normalized number of *mpeg1+* cells in the wound margin in vehicle- or nocodazole-treated conditions, *N =* 13 skin explants for vehicle conditions, *N =* 12 skin explants for nocodazole conditions. Mann Whitney U tests were used to determine significance in **(C-E),** two-way ANOVA followed by Bonferroni post-test was used to determine significance in **(F).** * = *p* < 0.05, ** = *p* < 0.01, *** = *p* < 0.001, **** = *p* < 0.0001. Timestamps denote mm:ss. Scale bars, 100 μm **(A, B).**

### Microtubule depolymerization alters actin distribution in migrating Langerhans cells

We next asked how microtubules promote efficient Langerhans cell migration to wounds. Previous works indicate multiple modes of crosstalk between the microtubule and actin cytoskeletons (Dogterom and Koenderink, 2019). One such mechanism involves sequestration of the RhoA activator GEF-H1/ARHGEF2 through binding to microtubules (Heck et al., 2012; Kashyap et al., 2019; Krendel et al., 2002). Microtubule depolymerization releases GEF-H1 from the microtubule lattice, leading to RhoA activation and subsequent changes in cell shape and actin distribution. Notably, our previous work suggests that a RhoA/Rho-associated kinase (ROCK)/myosin pathway promotes Langerhans cell migration to tissue wounds (Peterman et al., 2024). We hypothesized that Langerhans cell microtubule depolymerization alters RhoA activity and actin distribution, thereby disrupting migration towards wounds. To visualize F-actin in Langerhans cells and epidermal wounds, we used skin explants from *Tg(mpeg1.1:Lifeact-mRuby)* fish (Peterman et al., 2024). After explant, we pretreated scales with vehicle or nocodazole and scratched the epidermis. We followed individual cells as they migrated towards the wound and quantified Lifeact-mRuby ratios in the leading and trailing halves of migrating Langerhans cells. In vehicle-treated cells, we found roughly equal amounts of Lifeact-mRuby in the leading and trailing halves **(Figure 4A, A’, D)**. By contrast, nocodazole-treated cells contained an enrichment of Lifeact-mRuby in the trailing half, consistent with the idea depolymerizing microtubules increases RhoA activity at the rear of the cell **(Figure 4B, B’** (arrowheads)**, D)**. To inhibit a prominent downstream target of RhoA, we next used the ROCK inhibitor Y-27632 (Ishizaki et al., 2000), which we previously used to modulate Langerhans cell damage responses (Peterman et al., 2024). The addition of Y-27632 to nocodazole-treated explants restored Lifeact-mRuby ratios to near-control levels **(Figure 4C, C’, D)**. These data suggest that depolymerization of microtubules alters actin distribution through a ROCK-dependent mechanism in migrating cells and is one plausible cause for altered cell migration.

**Figure 4.**
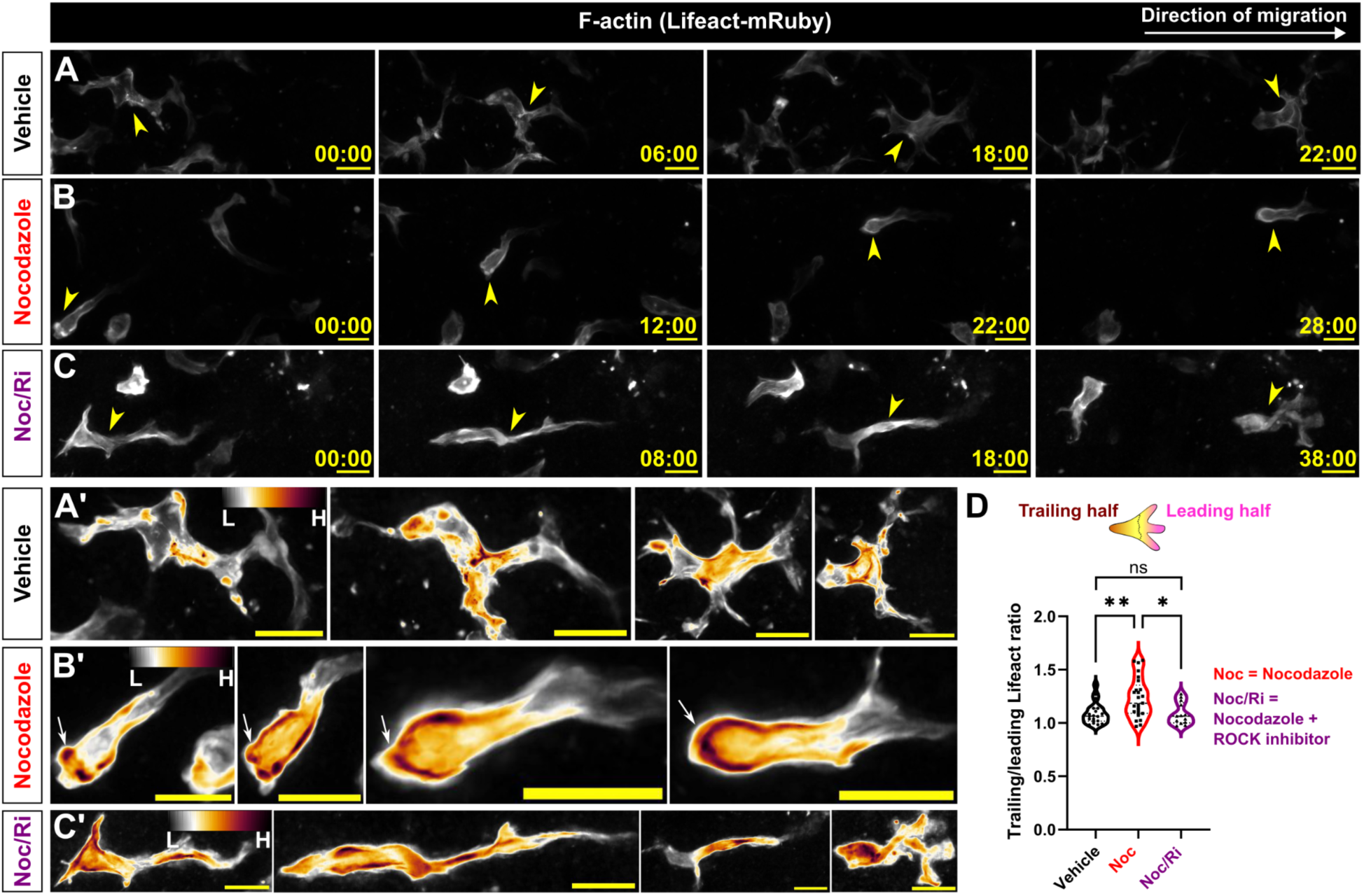
Microtubule depolymerization increases F-actin levels in a ROCK-dependent manner in the trailing halves of migrating Langerhans cells. **A-C.** Stills from time-lapse confocal microscopy showing migrating *Tg(mpeg1.1:Lifeact-mRuby)+* Langerhans cells treated with vehicle **(A)**, nocodazole **(B)**, or nocodazole+Y-27632 (Noc/Ri) **(C)**. Magnified insets in **(A’-C’)** are false-colored to show Lifeact-mRuby levels. Arrowheads in **(A-C)** indicate migrating cell, arrows in **(B’)** indicate increased Lifeact-mRuby levels in the trailing half of a nocodazole-treated cell. **D.** Violin plot quantifying the ratio of Lifeact-mRuby at the trailing and leading halves (see Materials and Methods for analysis detail), *n* = 20 cells tracked from *N* = 3 individual experiments in vehicle conditions, *n* = 25 cells tracked from *N* = 4 individual experiments in nocodazole-treated conditions, *n* = 13 cells tracked from *N* = 6 individual experiments in nocodazole+ROCK inhibitor-treated conditions. Significance was determined using one-way ANOVA followed by Bonferroni post-test. * = *p* < 0.05, ** = *p* <0.01. Timestamps denote mm:ss. Scale bars, 10 μm **(A-C)**.

### Pathfinding during directional cell migration involves MTOC decision-making

While imaging Langerhans cell migration to epidermal wounds, we observed that Langerhans cell bodies would often transiently pause while their dendrites continued to extend and retract **(Figure 5A-A’’’)**. Close examination of this phenomenon revealed that keratinocytes appeared to impede Langerhans cell migration to the wound.

**Figure 5.**
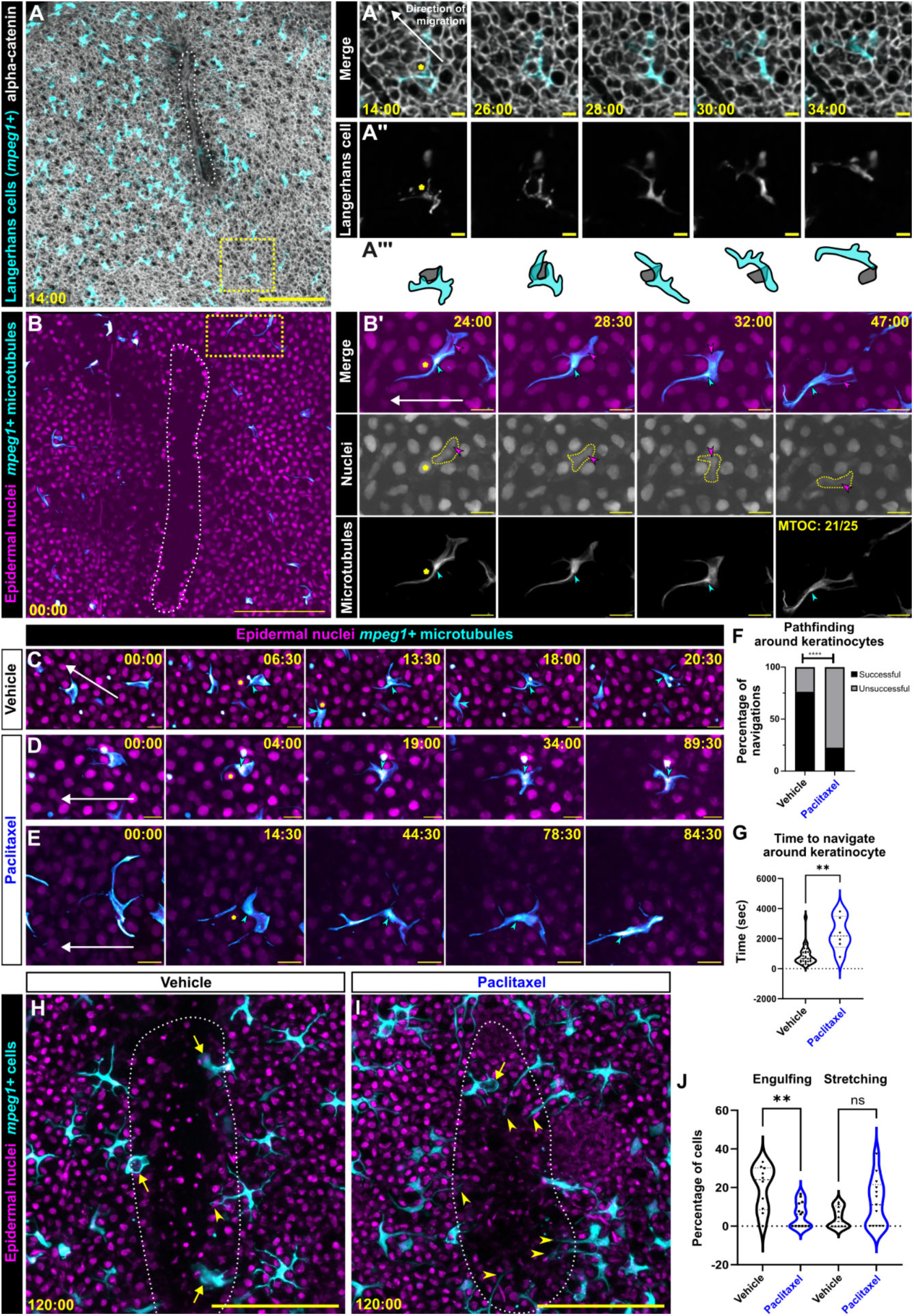
MTOC motility correlates with navigational pathfinding towards large epidermal wounds. **A.** Maximum intensity projection from time-lapse confocal microscopy of Tg(*mpeg1:mCherry)*;*Gt(ctnna1-Citrine)* skin explant. White dotted outline indicates epidermal scratch. Yellow box in **(A)** is magnified in **(A’,A’’)** insets as single *z*-slices. Yellow asterisk indicates obstacle keratinocyte. **B.** Representative still images from time-lapse confocal microscopy of *Tg(actb2:H2B-2x-mScarlet;mpeg1.1:EMTB-3xGFP)* skin explants. White dotted outline indicates epidermal scratch. Yellow box in **(B)** is magnified in **(B’)** insets. Yellow asterisk indicates obstacle nucleus, cyan arrowhead indicates MTOC, magenta arrowhead indicates Langerhans cell nucleus. 21 out of 25 (84%) of cells exhibit MTOC-first phenotype from *N* = 3 individual experiments. **C.** Representative still images from time-lapse confocal microscopy of vehicle-treated *Tg(actb2:H2B-2x-mScarlet;mpeg1.1:EMTB-3xGFP)*+ Langerhans cells navigating around obstacle keratinocytes towards a wound. **D, E.** Representative still images from time-lapse confocal microscopy of paclitaxel-treated *Tg(actb2:H2B-2x-mScarlet;mpeg1.1:EMTB-3xGFP)*+ Langerhans cells attempting to navigate around obstacle keratinocytes towards a wound. Yellow asterisks denote obstacle keratinocyte, cyan arrowhead denote MTOC. **F.** Quantification of navigation attempts in vehicle-treated and taxol-treated conditions, *n* = 38 attempts tracked from *N* = 3 individual experiments in vehicle conditions, *n* = 31 attempts tracked from *N* = 3 individual experiments in taxol conditions. **G.** Violin plots showing the length of time required to navigate around keratinocyte, *n* = 26 cells tracked from *N* = 3 individual experiments in vehicle conditions, *n* = 6 cells tracked from *N* = 3 individual experiments in taxol conditions. **H, I.** Representative still images from time-lapse confocal microscopy of *Tg(actb2:H2B-2x-mScarlet;mpeg1.1:YFP)* skin explants treated with vehicle **(H)** or paclitaxel **(I).** Yellow dotted line indicates wound, asterisks indicate phagocytic cells in wound margin, arrowheads indicate cells with stretched dendrites. **J.** Violin plots showing the percentage of cells exhibiting phagocytosis or stretching after 2 hours, *n =* 299 cells counted from *N =* 10 scales in vehicle conditions, *n =* 128 cells counted from *N =* 13 scales. Fisher’s exact test was used to determine significance in **(F)** by using the raw numbers of successful and unsuccessful navigations. Mann Whitney U test was used to determine significance in **(G)**. One-way ANOVA followed by Bonferroni post-test was used to determine significance in **(J).** ** = *p* < 0.01, **** = *p* < 0.0001. Timestamps denote mm:ss. Scale bars, 10 μm **(B’-E)**, 100 μm **(B, H, I)**.

A primary role for microtubules and the MTOC in immune cell function is dictating cell polarity and directionality of migration (Kopf et al., 2020; Kroll et al., 2023; Renkawitz et al., 2019). This is accomplished by positioning microtubule-associated proteins including vesicle cargo, motors and focal adhesions. Certain cell types, such as dendritic cells, often exhibit a nucleus-first mode of migration, while neutrophils can exhibit either MTOC- or nucleus-first migration (Etienne-Manneville, 2013). Therefore, we first asked how the MTOC was positioned during directional migration and whether or not its position corresponded to pathfinding decisions. To facilitate imaging nuclear positioning of all epidermal cells, we generated a transgenic line in which the quasi-ubiquitous *beta-actin2* (*actb2*) promoter (Kobayashi et al., 2014) drives expression of the histone H2B fused to a tandem repeat of the red fluorescent protein mScarlet (Bindels et al., 2017) [*Tg(actb2:H2B-2x-mScarlet)*]. By scratching skin explants from *Tg(mpeg1.1:EMTB-3xGFP;actb2:H2B-2x-mScarlet)* fish, we could track the Langerhans cell MTOC and nucleus as the cell migrated towards the wound **(Figure 5B; Supplemental Video 7)**. As Langerhans cells migrated towards the wound, we observed that cells would often encounter surrounding nuclei and pause **(Figure 5B’)**. While paused, Langerhans cell dendrites explored the space beyond the keratinocyte obstacle. In 21/25 instances, the MTOC preceded the nucleus past the obstacle keratinocyte **(Figure 5B’)**.

Prior studies found that paclitaxel-mediated microtubule stabilization can inhibit cell migration (Ganguly et al., 2011; Wang et al., 2019) and centriole motility (Ching et al., 2022). To determine the effects of microtubule stabilization on Langerhans cell migration, we performed the scratch injury assay in the presence of paclitaxel or vehicle control. Using skin explants from *Tg(mpeg1.1:EMTB-3xGFP;actb2:H2B-2x-mScarlet)* fish and observing MTOC position as a proxy for successful navigation, we examined how cells and microtubules behaved after paclitaxel treatment. Vehicle-treated Langerhans cells successfully navigated past obstacle cells 76% of the time with an average navigation time of 675 s **(Figure 5C,F,G; Supplemental Video 8)**. By contrast, paclitaxel-treated Langerhans cells failed obstacle navigation 77% of the time **(Figure 5D-F)**. When paclitaxel-treated cells managed to successfully navigate past their obstacle, navigation times were significantly longer compared to control cells, averaging 2175 s **(Figure 5G)**. To observe possible outcomes from this inability to navigate the epidermis, we tracked Langerhans cell behaviors as they attempted to enter the wound margin using skin explants from *Tg(mpeg1.1:YFP;actb2:H2B-2x-mScarlet)* fish. After 2 hours in control conditions, we observed a subset of Langerhans cells had entered the closing wound and engulfed debris (**Figure 5H**, arrows, **Figure 5J**, average of 20% of cells per wound margin ROI). Less frequently, we noted Langerhans cells on the wound margin did not enter the recovering wound but instead projected an extended dendrite into the region (**Figure 5H**, arrowhead, **Figure 5J**, average of 4.7% of cells per wound margin ROI). In paclitaxel-treated conditions, Langerhans cells entered the wound margin to engulf debris at a significantly lower rate, and we observed a moderate increase in the number of cells that probed the wound with extended dendrites **(Figure 5H**, arrowheads, **Figure 5J)**. Together, these results suggest that Langerhans cells use MTOC positioning to aid in circumventing cellular obstacles during migration and that stabilizing microtubules inhibits navigational pathfinding to wounds.

### Summary: Microtubules dictate immune responses to tissue damage

How immune cells dynamically and rapidly navigate densely packed epithelia to respond to injury remains largely unknown. Here, we use scale explants from adult zebrafish skin as a tractable model to examine tissue-resident macrophage behaviors within a native epithelial microenvironment and apply acute chemical perturbations to modulate the microtubule cytoskeleton. We show that microtubule depolymerization alters Langerhans cell morphology, including dendrite number and length. We find that the MTOC rapidly moves towards the phagosome during debris engulfment and that microtubules are partially required for efficient debris engulfment. What functions might the microtubule cytoskeleton play during phagocytosis? Using cultured macrophages, (2007; Athlin et al., 1986; Walter et al., 1980) found that the MTOC reorients during phagocytosis of antibody-coated beads and red blood cells and that microtubules were strictly required for engulfment. The authors posited that the MTOC can increase interactions between the Golgi apparatus and the phagosome, which may facilitate subsequent cross-presentation events between the phagocyte and other immune cells (Eng et al., 2007). By contrast, Möller et al., (2022) found that depolymerization of microtubules changed how zebrafish microglia encounter and engulf apoptotic corpses *in vivo*. In the absence of microtubules, microglia relied more heavily on a motility-based engulfment strategy (Möller et al., 2022), similar to our own observations **(Figure 2H-J)**, highlighting the flexible engulfment abilities of phagocytes within native microenvironments.

During directed migration to scratch wounds, we found that Langerhans cells frequently paused at sites of keratinocyte contact, which present potential decision points, akin to microfluidic mazes developed to study migration decisions *in vitro* (Ambravaneswaran et al., 2010; Kopf et al., 2020; Kroll et al., 2023; Renkawitz et al., 2018; Tweedy et al., 2020). By imaging Langerhans cell microtubules, we found that MTOC positioning frequently, but not exclusively, correlated with pathfinding during migration. Perturbing microtubule polymerization and stability affected migration efficiency and actin localization. Our data suggest two non-mutually exclusive models in which microtubules and the MTOC promote efficient migration: through the sequestration and release of actin-regulating proteins, and through decision making in directional migration around obstacles. Further dissecting these mechanisms will require developing methodology to specifically manipulate actin-microtubule crosstalk and the MTOC in Langerhans cells.

Collectively, our work demonstrates requirements for microtubules in steady-state organ surveillance and directed cell migration by tissue-resident macrophages during wound repair. Uncovering the details and regulation of Langerhans cell motility are important not only for understanding responses to tissue injury, but also for how these cells initially seed the skin, and emigrate from the skin to secondary immune organs.

## Acknowledgements

We thank the LSB Aquatics staff for animal care, and Dan Fong and Wai Pang Chan for imaging support. We thank Anna Huttenlocher’s lab for providing the *mpeg1.1:EMTB-3xGFP* plasmid. EB3-GFP was a gift from Anna Akhmanova (Addgene plasmid #190164; http://n2t.net/addgene:190164; RRID:Addgene_190164).

## Author contributions

Conceptualization: E.P., J.P.R.; Methodology: E.P.; Formal analysis: E.P.; Investigation: E.P.; Resources: E.P., A.M, I.A.S; Writing-original draft: E.P.; Writing-review & editing: E.P., J.P.R.; Visualization: E.P., J.P.R.; Supervision: E.P., S.G.M, J.P.R.; Project administration: E.P., J.P.R.; Funding acquisition: E.P., J.P.R.

## MATERIALS AND METHODS

### Key resources table

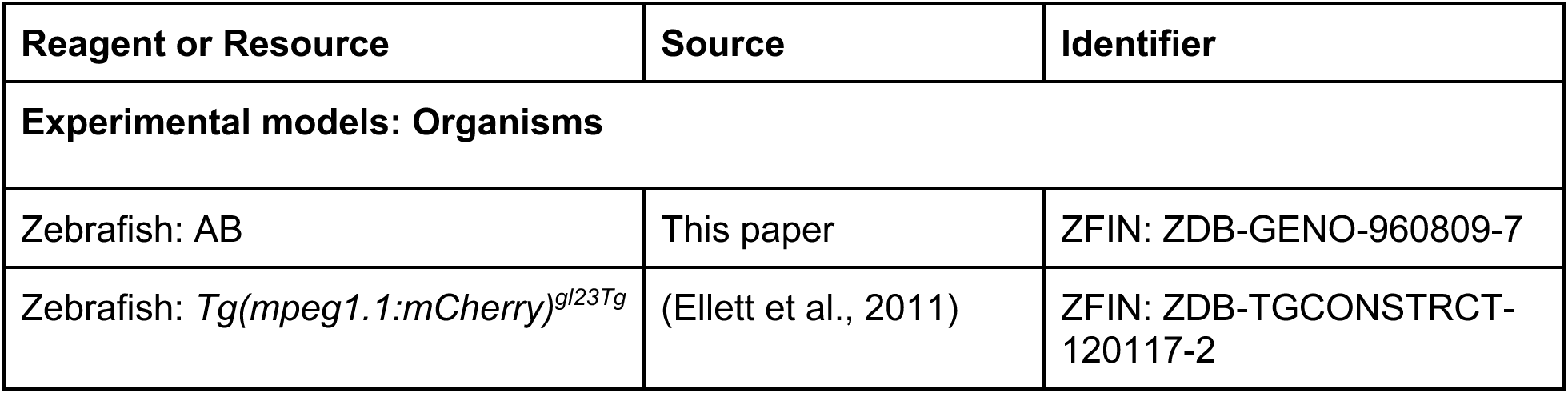
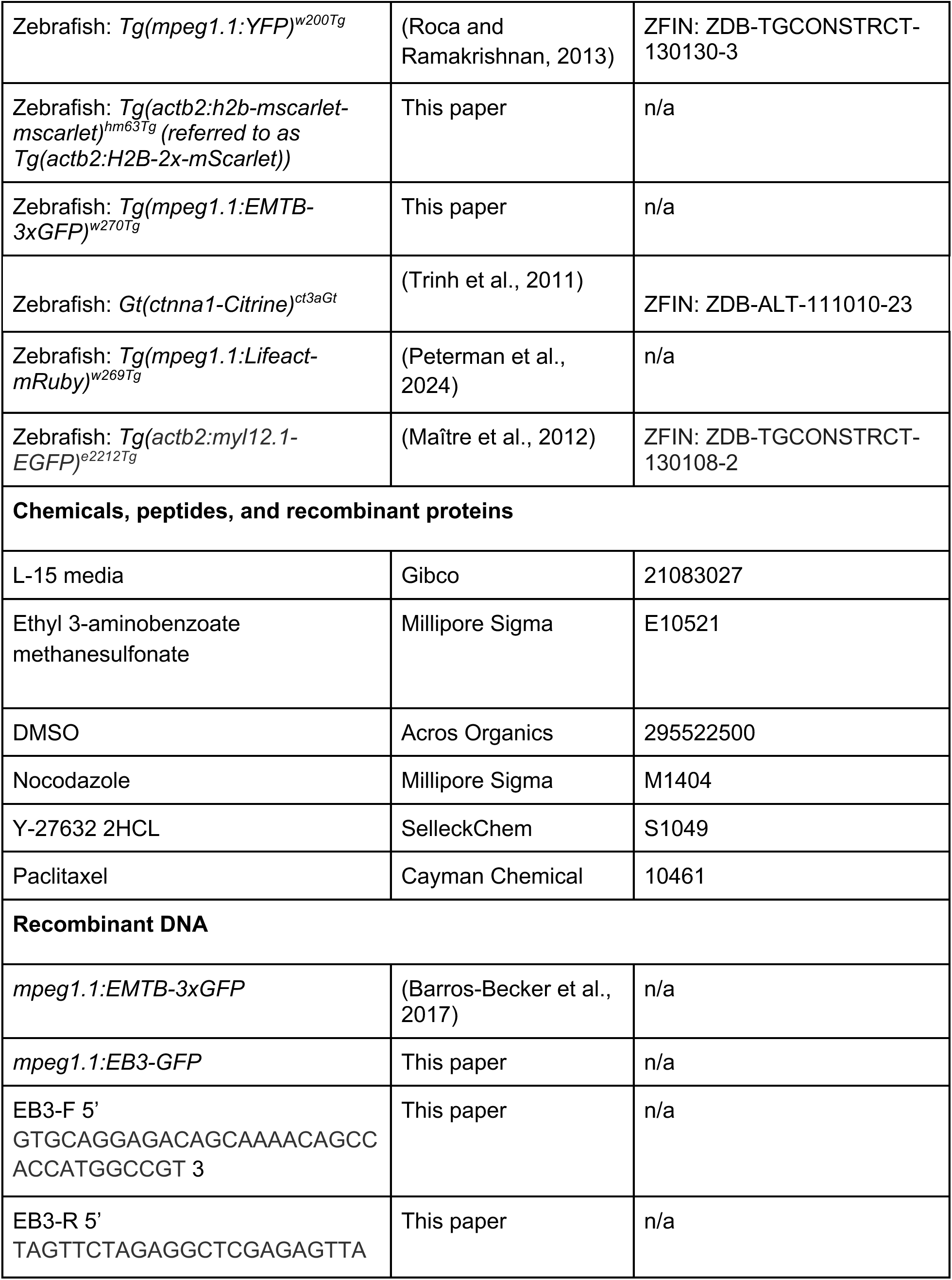
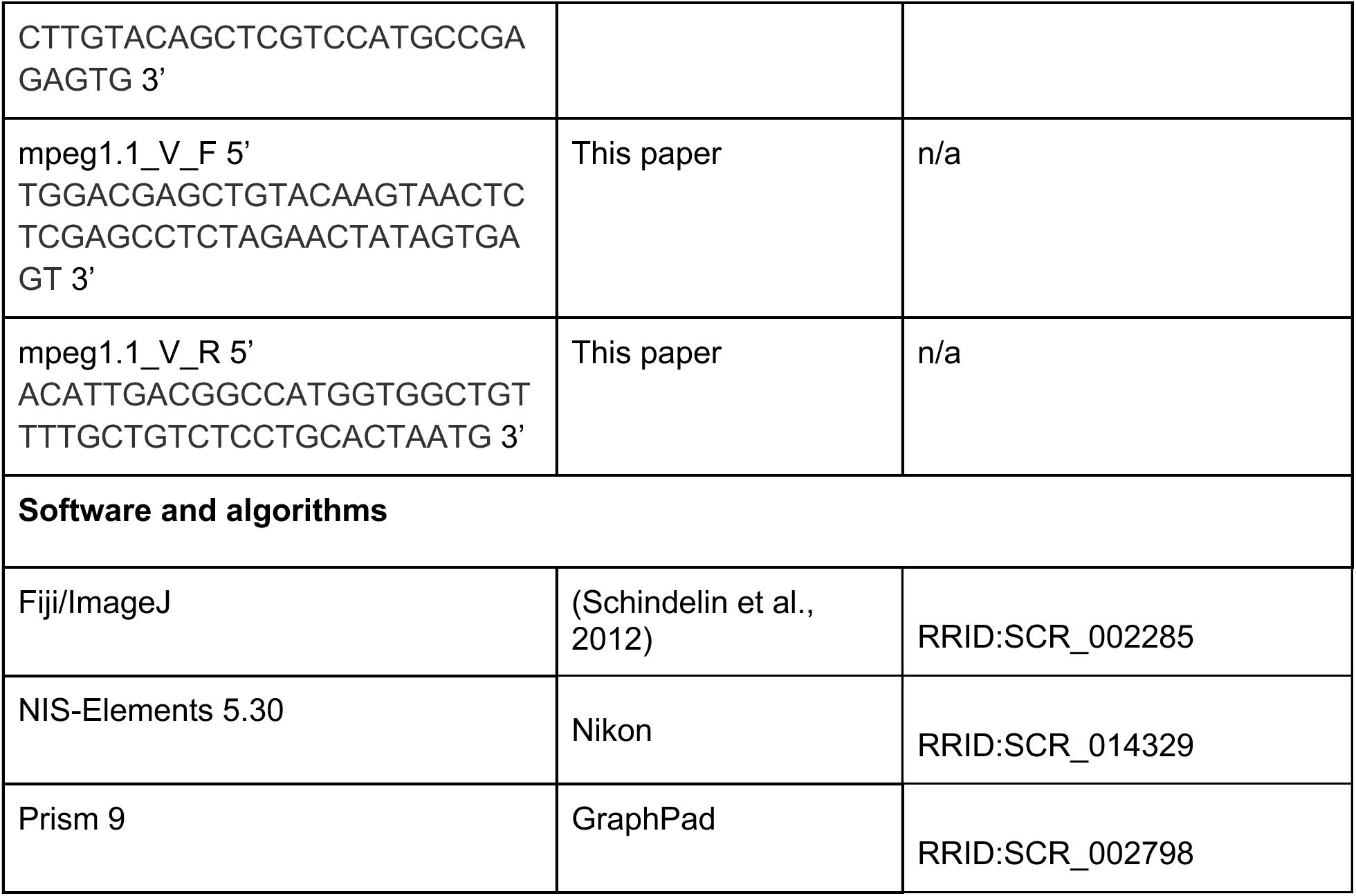

### Zebrafish husbandry

Zebrafish were housed at 26-27°C on a 14/10 h light cycle. The strains used are listed in the Key Resources Table. Animals aged 6-18 months of either sex were used in this study. All zebrafish experiments were approved by the Institutional Animal Care and Use Committee at the University of Washington (Protocol #4439-01).

### Generation of transgenic zebrafish

To generate *Tg(mpeg1.1:EMTB-3xGFP)^w270Tg^,* a previously published *mpeg1.1:EMTB-3xGFP* plasmid (Barros-Becker et al., 2017) and *tol2* mRNA were injected into AB embryos at the 1-cell stage, and embryos were raised to adulthood. Adults were screened for GFP+ cells in the skin, and GFP+ adults were then outcrossed to wild-type partners. F1 fish were raised to adulthood, where GFP expression was assessed in Langerhans cells to identify a line to propagate for this study.

To generate *Tg(mpeg1.1:EB3-GFP)* G0 animals, our published *tol2* plasmid containing *tol2* sites, the *mpeg1.1* promoter and the *cryaa:DsRed* transgenesis marker (Peterman et al., 2024) was modified to insert EB3-GFP downstream of the *mpeg1.1* promoter. To amplify EB3-GFP, the Addgene plasmid #190164 was used as template and amplified with the primers EB3-F and EB3-R. Our previously published *mpeg1.1* promoter-containing plasmid was amplified using the primers mpeg1.1_V_F’ and mpeg1.1_V_R. Gibson assembly was used to construct the final pTol2-*mpeg1.1:EB3-GFP;cryaa:DsRed* plasmid. *tol2* mRNA and pTol2-*mpeg1.1:EB3-GFP;cryaa:DsRed* plasmid DNA were injected into *Tg(mpeg1.1:mCherry)* embryos at the 1-cell stage and larvae were screened for *cryaa:DsRed* expression. Positive animals were raised to adulthood, where GFP expression was assessed in Langerhans cells.

To generate *Tg(actb2:h2b-mscarlet-mscarlet)^hm63Tg^* animals, Gibson assembly was used to create an H2B-2x-mScarlet fusion protein by adding a linker between two copies of mScarlet and fusing this to the C-terminus of human histone 2B. This was cloned into the pMTB expression construct that contains the zebrafish *beta-actin 2* (*actb2*) enhancer/promoter inside mini-Tol2 transposon system (Xiong et al., 2013), which was used for generating stable transgenics. *tol2* mRNA and plasmid DNA were injected into AB embryos at the 1-cell stage. Potential founders were screened for bright red ubiquitous fluorescence and outcrossed to obtain a stable line.

### Scale removal

For scale removal, adult fish were anesthetized in system water containing 200 µg/ml buffered tricaine. Individual scales were removed with forceps and placed onto 6 mm plastic dishes, epidermis side up, and allowed to adhere for 30 seconds before adding L-15 medium prewarmed to room temperature (22°C). Following scale removal, animals were recovered in system water.

### Scale injury assay

For the epidermal scratch assay (Figures 3, 4, 5), one pair of forceps was used to assist in pinning the scale down by contacting a region devoid of epidermis on the anterior scale surface. A second pair introduced the scratch in the middle of the epidermis. Scratches were only used for data collection if they did not extend to the edge of the scale and were oval in shape (as shown by representative images in Figure 3A, B). When treating scales with pharmacological agents, scales were removed, placed in L-15, and pretreated with vehicle control or pharmacological agent. Nocodazole-treated (50 µM) scales were pretreated for 10 minutes, nocodoazole- (50 µM) + ROCKi- (25 µM) treated scales were first treated with Y-27632 (40 minutes) then nocodazole (10 minutes), and paclitaxel-treated (25 µM) scales were pretreated for 30 minutes. After preincubation, scales were placed under a dissecting microscope, scratched, and placed under the confocal microscope for imaging.

### Microscopy and live imaging

An upright Nikon Ni-E microscope equipped with an A1R MP+ confocal scanner, a piezo *z*-drive (Mad City Labs Nano-F450), and a 25× water dipping objective (1.1 NA; Nikon MRD77220) was used for all experiments. Laser powers between 0.5 and 1% were used for 488 and 561 nm lasers in conjunction with 521/42 and 600/45 emission filter sets for GFP and RFP channels, respectively. *Z*-stacks were acquired using a resonant scanner and post-processed to remove noise using the denoise.ai function in NIS-Elements.

### Steady-state chemical treatments

For nocodazole and paclitaxel treatments, scales were removed, placed onto dry 6 mm dishes, and incubated in 5 ml L-15 media. Imaging commenced for at least 10 minutes before careful addition of chemicals while on the microscope stage. Appropriate vehicle controls (DMSO) were used at equivalent %v/v.

### Laser-induced cell damage

For laser-induced cell damage, scales were mounted as described above. Target cells at least 1 cell distance away from a Langerhans cell (∼5-15 microns) and within the same *z*-plane were located and ablated using a UGA-42 Caliburn pulsed 532 nm laser (Rapp OptoElectronic). The laser was focused through a 25× objective at 4× zoom. Ablation was produced in the focal plane using 15-20% power at a single point within a nucleus, firing 3 times for 3 seconds each using a custom NIS-Elements macro. In the presence of nocodazole (Figure 2), scales were first pretreated with nocodazole (50 µM) for 10 minutes, followed by laser-induced cell damage and subsequent confocal imaging.

### Image analysis

#### Dendrite metrics

Dendrites were counted as microtubule positive if the ratio of EMTB-3xGFP/mCherry in the dendrite was greater than 0.5. To quantify dendrite number and length in Figure 1, dendrites were manually counted and measured every 10 minutes throughout the treatment time course. Numbers and lengths across the time course were averaged to obtain values shown in Figure 1.

#### Sholl analysis

To calculate Sholl crossings, 5 timepoints from a time-lapse micrograph were selected for each cell. Each cell was thresholded and the Sholl Analysis Function in the Neuroanatomy plugin in ImageJ was used to calculate Sholl crossings (Arshadi et al., 2021; Ferreira et al., 2014). The resulting visualization was manually inspected to ensure accuracy. The sum of the radii crossings for each cell across the time points was calculated, and the percentage of total crossings was summed for each condition.

#### Cell motility

To calculate cell motility metrics (displacement, total distance traveled, meandering index) in Figure 3, cells were thresholded and tracked using the LAP algorithm in ImageJ (Ershov et al., 2022; Tinevez et al., 2017). Cells were only tracked and quantified if they could be tracked throughout the entire time course. Cells were excluded if their displacement was less than 10 microns. To quantify the meandering index for each individual cell, the displacement of that cell was divided by its total distance traveled.

To calculate the cell number in wound margin in Figure 3F, a rectangular region of 150 microns wide and (length of wound x 1.2) microns long was drawn around the wound. The number of cells was counted for each timepoint and normalized to time 0. Cells were only counted if >50% of their cell body was within the ROI. To calculate the engulfing and stretching phenotypes in Figure 5, the total number of cells at 2 hours within the wound margin (rectangular region of 150 microns wide and (length of wound x 1.2) microns long) was counted. A second region was drawn around the closing wound (as seen in Figure 5H, I). Cells were scored as engulfing if they were within the wound and presenting a rounded, phagocytic phenotype. Cells were scored as stretching if they extended dendrites, but not the cell body, into the wound. Frames prior to the 2 hour time were used as references to score if cells were morphologically changing into either of these phenotypes.

#### Trailing and leading edge Lifeact intensity

To calculate the trailing/leading edge Lifeact-mRuby ratio in Figure 4, a set of custom ImageJ macros were used. Migrating cells were thresholded and outlined over time. The outlines were used as ROIs, bisected to make a trailing half and leading half, and mean Lifeact-mRuby intensity in each half was quantified at each time point. The mean of means for trailing and leading halves over the course of migration was calculated and the ratio of trailing/leading was plotted.

#### MTOC motility and positioning

MTOCs were manually tracked in ImageJ in order to quantify their speed and distance from phagosome (Figure 2A-G). To score MTOC decision making in Figure 5B, a decision was scored as MTOC-first if the MTOC was the first structure to move past the obstacle nucleus. A decision was scored nucleus-first if the nucleus preceded the MTOC. To score successful navigation and navigation times in Figure 5, the difference in time between an MTOC first encountering a nucleus and its successful progression past the obstacle was calculated. Navigations were scored unsuccessful if the MTOC did not progress past the obstacle nucleus within at least a 55 minute observation window.

### Statistical analysis

GraphPad Prism was used to generate graphs and perform statistical analyses. At least three individual biological experiments were performed unless otherwise noted. Tests used and number scales or cells/ROIs are described in each figure legend.

## SUPPLEMENTAL FIGURES

**Supplemental Figure 1.**
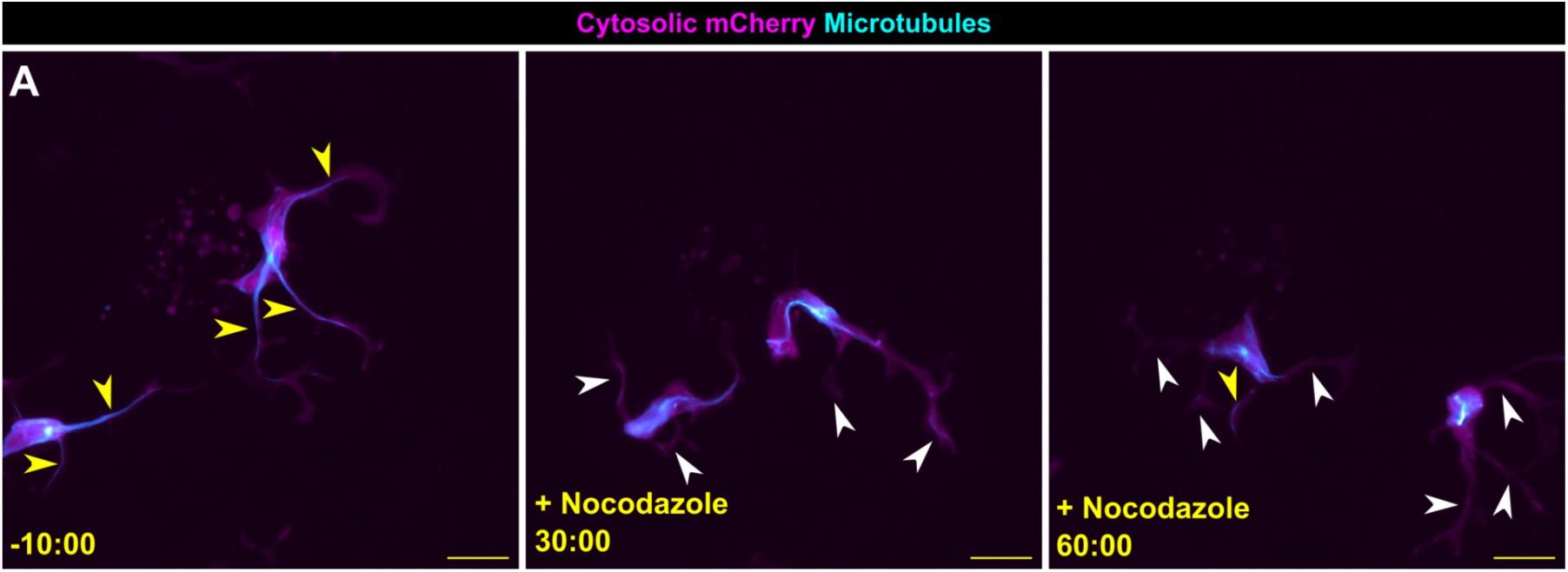
Confirmation of the EMTB-3xGFP reporter for tracking microtubules. **A.** Representative images from confocal time lapse microscopy of *Tg(mpeg1.1:mCherry;mpeg1.1:EMTB-3xGFP)*+ Langerhans cells showing effects of nocodazole treatment on EMTB signal within Langerhans cells. Yellow arrowheads indicate EMTB-positive dendrites, white arrowheads indicate EMTB-negative dendrites. Timestamps denote mm:ss relative to the addition of nocodazole. Scale bar, 10 μm.

## SUPPLEMENTAL VIDEO LEGENDS

**Supplemental Video 1.** Time-lapse microscopy of Langerhans cell (magenta; *Tg(mpeg1.1:mCherry)*) expressing the microtubule reporter EMTB-3xGFP (cyan; *Tg(mpeg1.1:EMTB-3xGFP)*). Scale bar, 10 μm.

**Supplemental Video 2.** Time-lapse microscopy of Langerhans cell (magenta; *Tg(mpeg1.1:mCherry)*) expressing the plus-end microtubule reporter EB3-GFP (cyan). Scale bar, 10 μm.

**Supplemental Video 3.** Time-lapse microscopy of Langerhans cells (white; *Tg(mpeg1.1:YFP)*) treated with vehicle, nocodazole, or paclitaxel in steady-state conditions. Scale bar, 20 μm.

**Supplemental Video 4.** Time-lapse microscopy of a Langerhans cell (magenta; *Tg(mpeg1.1:mCherry)*) MTOC during debris engulfing via tracking of EMTB-3xGFP foci (*Tg(mpeg1.1:EMTB-3xGFP)*). Asterisk indicates site of keratinocyte laser ablation, white line traces the MTOC motility during engulfment. Scale bar, 10 μm.

**Supplemental Video 5.** Time-lapse microscopy of Langerhans cell (white; *Tg(mpeg1.1:YFP)*) engulfing cellular debris generated after laser-induced damage of keratinocytes. Cells are treated with vehicle or nocodazole. Asterisk indicates site of keratinocyte laser ablation. Scale bar, 10 μm.

**Supplemental Video 6.** Time-lapse microscopy of Langerhans cells (cyan; *Tg(mpeg1.1:mCherry)*) reacting to epidermal wounds (epidermal cells labeled in white; *Gt(ctnna1-Citrine)*). Cells are treated with vehicle or nocodazole. Scale bar, 100 μm.

**Supplemental Video 7.** Time-lapse microscopy of MTOC-labeled Langerhans cell (cyan; *Tg(mpeg1.1:EMTB-3xGFP)*) navigating around obstacle nuclei (magenta;*Tg(actb2:H2B-2x-mScarlet)*) in order to reach the epidermal wound. Asterisk indicates an obstacle nucleus. Scale bar, 10 μm.

**Supplemental Video 8.** Time-lapse microscopy of MTOC-labeled Langerhans cell (cyan;*Tg(mpeg1.1:EMTB-3xGFP)*) navigating around obstacle nuclei (magenta;*Tg(actb2:H2B-2x-mScarlet)*) in order to reach the epidermal wound. Cells are treated with vehicle or paclitaxel. Asterisk indicates an obstacle nucleus. Scale bar, 10 μm.

