## Supplementary figures and images for "Microtubule-dependent cell polarity regulates skin-resident macrophage phagocytosis and directed cell migration"

### Supplemental Figure 1

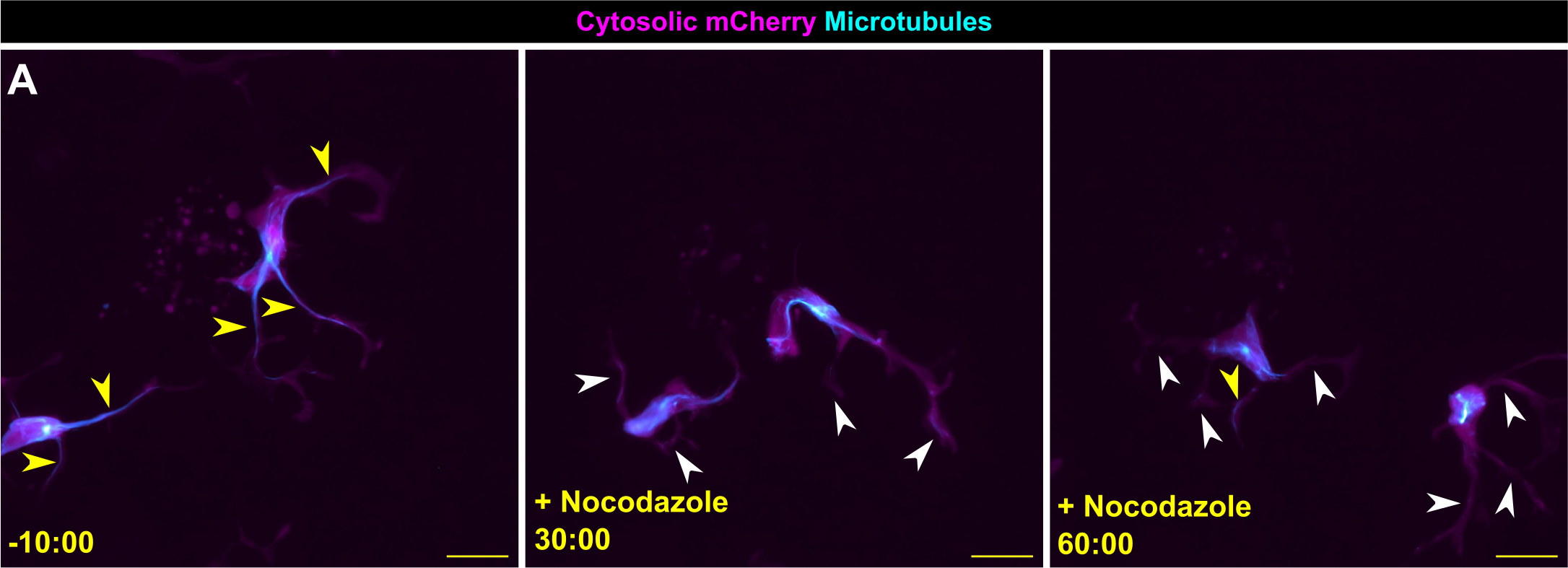
